# Harnessing Contextual Embeddings: A Deep Learning Framework for Predicting PCR Amplification Using BERT Tokenization

**DOI:** 10.1101/2024.11.23.624986

**Authors:** Niloofar Latifian, Naghme Nazer, Amir Masoud Jafarpisheh, Babak Khalaj

## Abstract

Polymerase Chain Reaction (PCR) is a widely used molecular biology technique to amplify DNA sequences. PCR amplification is affected by factors such as binding dynamics and primer-template interactions. This study aims to reduce the time and cost of the experiment by predicting PCR outcomes based on these factors. To achieve this, we first identify the most stable binding sites for each primer-template pair by calculating the Gibbs free energy. Then, we propose a unique labelling strategy that captures primer-template interactions in the binding sites by analyzing match and mismatch positions. We categorize a set of English words into two semantically distinct groups: one for match positions and another for mismatch positions. Words within each group have a higher cosine similarity to one another than to words in the opposing group. We assign the corresponding word to each base pair based on whether it is a match or a mismatch. The labelled sequence is then tokenized with BERT, serving as input to a CNN-BiLSTM model. Achieving 85.9% accuracy, this approach outperforms prior methods and pioneers BERT-based analysis in primer-template binding. Crucially, the model demonstrates significantly better sensitivity and overall AUC compared to prior work, while maintaining highly competitive specificity. This indicates a far more balanced and robust capability to correctly distinguish both successful and failed PCR outcomes, which is vital for reliable experimental prediction.

**Highlights:**

- Selecting the most important features for PCR amplification using Random Forest Classifier
- Proposing a new labelling approach to represent the matches and mismatches between PCR primers and templates
- Using BERT tokenizer to tokenize the corresponding representation of matches and mismatches
- Augmenting the data based on the semantic similarities of the words in the BERT tokenizer
- Using CNN-BiLSTM to predict PCR amplification results

## 1. Introduction

Polymerase Chain Reaction (PCR) is a widely utilized molecular biology technique that enables the exponential amplification of specific DNA sequences. It allows researchers to generate millions or even billions of copies of a target DNA fragment from a minimal starting sample [1]. The amplification of a specific segment of DNA enables its analysis in various studies. One of the recently popular and well-known applications of this process is the detection of the coronavirus in patients [2]. The PCR process begins with extracting a DNA template containing the target sequence, which can be sourced from cells, tissues, or environmental samples. Once the template is prepared, short DNA sequences called primers are designed and synthesized to bind to regions flanking the target sequence. The primers that bind to the antisense strand (3’ to 5’) and sense strand (5’ to 3’) are named forward primer and reverse primer, respectively. These primers act as initiation sites for DNA synthesis. The design of specific and efficient primers is crucial to ensuring the successful amplification of the target sequence, as they must precisely complement both ends of the DNA fragment to be amplified [3].

In a laboratory, several factors critically impact the success of PCR amplification. Parameters such as primer sequence, template DNA concentration, annealing temperature, and cycle conditions must be carefully optimized for each experiment. Important considerations in primer design include Gibbs free energy (Δ*G*) for stable primer-template binding [4], avoiding 3’ mismatches to ensure proper extension [5], [6], and maintaining a balanced GC content (ideally 40-60%) [7] to promote stable hybridization. Additionally, secondary structures like hairpins or dimers must be minimized to ensure proper binding [8].

Molecular biologists currently rely on established software tools like Primer3 [9], OLIGO7 [10], and PrimerSelect [11] to design primers based on these heuristics. While these tools are essential for the initial design, they are ultimately unreliable at predicting the final, binary outcome (amplified or not amplified) of the experiment. A significant percentage of computationally designed primer pairs fail in the lab, forcing researchers into costly, time-consuming cycles of redesign, re-synthesis, and re-testing. This failure rate is especially problematic and expensive in high-throughput applications such as large-scale genotyping or mutation screening, multiplex PCR (simultaneously amplifying dozens of targets), and preparation of hundreds of samples for next-generation sequencing (NGS) library preparation. In these scenarios, the ability to accurately predict failure before committing to the experiment offers enormous value by de-risking the entire workflow, saving significant time, reagents, and money, and dramatically increasing overall throughput [12].

Some prior works tried to optimize the PCR amplification process by selecting better primers [13, 14, 8] or improving its efficiency [15, 16]. In [13, 14], a teaching-learning-based method for selecting the best primer to bind to the template was proposed. This approach employs iterative algorithms that mimic teaching and learning dynamics to refine primer selection based on multiple criteria, potentially leading to more effective and specific amplification. In [8], an algorithm was developed to prioritize the similarity between designed primers and their target sequences. By focusing on this crucial aspect, the algorithm aims to enhance specificity and reduce off-target amplification. In [15], an Artificial Neural Network was designed with nine experimental inputs of PCR experiment to estimate its efficiency. This approach allows for more accurate predictions of amplification success based on input conditions. In [16], a web tool was designed to identify some parameters that affect PCR efficiency using a statistical approach. This platform enables researchers to systematically evaluate the impact of various factors on amplification outcomes.

Despite the extensive application of PCR in molecular biology, relatively few studies have focused specifically on predicting the outcome of PCR amplification. Cordaro et al. utilized a random forest classifier to Predict PCR results for a diverse set of templates ranging from human, plant, monkey, bacteria, and plasmid DNA [12]. This study included 290 PCR reactions consisting of 109 unique primer pairs. They considered 15 different features of the experiment to train their model, which yielded 81% accuracy. Kronenberger et al. [17] also used a random forest classifier to predict PCR amplification results based on 10 parameters affecting the experiment. The study focused on environmental DNA (eDNA) and paired 10 qPCR assays with 82 synthetic gene fragments. They conducted 530 specificity tests, achieving accuracies of 92.4% with SYBR Green and 96.5% with TaqMan hydrolysis probes. Döring et al. presented a dataset of 47 immunoglobulin heavy chain variable sequences and 20 primers [18]. They developed a logistic regression model considering 22 parameters for predicting the amplification status with an Area Under the Curve (AUC) of 95%. Kayama et al. expressed each base pair in the primer-template sequence as a five-lettered word and then predicted the PCR results using an RNN model based on the modeled sequence [19]. They reached 80% accuracy on a dataset of 72 PCR primers capable of amplifying 31 DNA templates. Bai et al. predicted PCR amplification results using the same dataset and same modeling approach, with a CNN BiLSTM gaining 82% accuracy [20].

While numerous tools have been developed for primer design, predictive models that comprehensively account for the diverse factors influencing PCR efficiency—such as primer-template interactions, thermodynamic constraints, reaction conditions, and the potential for non-specific binding—remain limited. Many of the existing approaches suffer from low predictive accuracy, lack robustness across varying experimental conditions, or require manual tuning of experimental parameters. This highlights a pressing need for more systematic and accurate methods to predict PCR outcomes, particularly as PCR continues to be a foundational technique in molecular biology and genomics.

Recently, transformer-based models such as DNABERT [21], DNABERT-2 [22], and HyenaDNA [23] have shown strong performance in modeling genomic sequences. DNABERT and DNABERT-2 leverage k-mer tokenization combined with BERT-style embeddings to capture contextual dependencies in DNA sequences, while HyenaDNA introduces an attention-free architecture capable of learning from base-pair resolution input over extremely long ranges. Despite their success in tasks such as promoter prediction and chromatin profiling, these models have not been applied to modeling the specific primer-template interactions central to PCR amplification. This gap reveals the need for a specialized sequence representation and modeling approach that can effectively capture the biochemical and positional characteristics of primer binding, thereby improving the prediction of PCR outcome.

In this study, we introduce a novel, accurate, and robust predictor for PCR amplification outcome. The intended use case for this tool is as a final filter after traditional design software (e.g., Primer3) has generated a list of candidate primers. The researcher submits the candidates to our predictor to de-risk the success rate before purchasing expensive synthesis. The model is also ideal for in silico screening of large libraries of existing primer pairs, where experimental validation is infeasible due to cost or time constraints. By providing a reliable and accurate prediction, our method simplifies experimental workflows, minimizes the reliance on repetitive trial-and-error optimization, and increases the overall throughput and reliability of PCR-based research.

In our approach, we predict the amplification result using a multi-input, multi-branch architecture. The primary branch processes the primer-template sequences as shown in Figure 1. First, we use a set of primers and templates and their amplification results as input data, and then employ a feature selection method to select the most important features in predicting the outcome. On the basis of the results of feature selection, we need to identify the binding site using the Gibbs free energy. Then, we label each match and mismatch in the primer-template combination using English words and do a data augmentation for sequences. After that, we utilize a BERT tokenizer [24] to tokenize the modeled sequence. Then, we process the sequence by generating contextual embeddings from a fine-tuned BERT model, and then passing through a Convolutional Neural Network (CNN) and a Bidirectional LSTM (BiLSTM) to capture complex local and sequential patterns. In parallel, a second branch of our model processes pre-calculated biological features (like GC content and mismatch counts) through a simple dense network. Finally, the feature vectors from branches are concatenated and passed through a dense classifier to produce the final prediction. To the best of our knowledge, this is the first study to use a BERT tokenizer for modeling interactions between base pairs. Our method aims to simplify experimental workflows and improve the efficiency of PCR-based applications.

**Figure 1:**
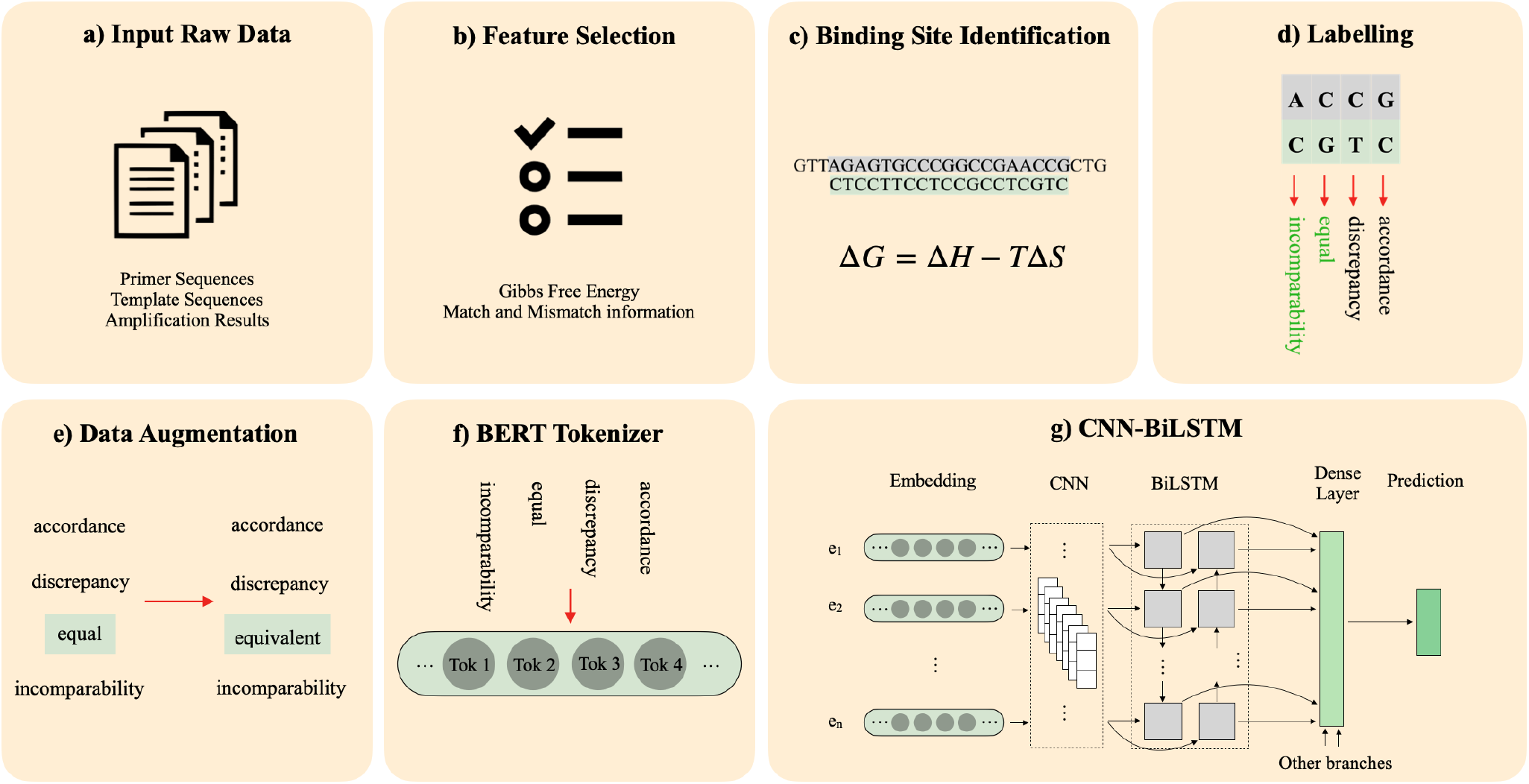
Overview of the primary branch of the pipeline for PCR amplification prediction. (a) Input data from a dataset of primer-template pairs with known amplification outcomes (b) Feature selection from experimental data using an embedded random forest method. Gibbs free energy and some information about matches and mismatches are the most important features. (c) Identification of primer binding sites on the template by calculating and finding the binding site with the minimum Gibbs free energy across potential binding positions. (d) Detection and labelling of match/mismatch positions within and around the binding sites, encoded using semantically related English words. (e) Data augmentation by duplicating sequences and replacing some original labels with semantically similar English words to increase diversity and model robustness. (f) Tokenization of encoded sequences using a BERT tokenizer to capture contextual patterns before deep learning-based prediction. (g) Classification of sequences using an CNN-BiLSTM model to learn contextual dependencies and highlight informative positions. the output will then served as the input to a dense layer with combination of two other branches of our model

## 2. Materials and Methods

### 2.1. Datasets

To predict the outcomes of PCR, it is essential to first collect appropriate data. This involves conducting and recording the results of the PCR process on a set of primers and DNA samples, categorizing the outcomes as either “amplified” or “not amplified.” There are few datasets in the scientific literature that document experimental features and their final results in this manner, particularly those that include “not amplified” samples. In this study, we utilized datasets provided by Döring et al. [18], and Kayama et al. [19], that contain both desired outcomes (Figure 1-a).

The dataset of Döring et al. contains the amplification status of 47 IGHV genes and 20 primers. Their study includes information about 22 different features related to PCR. In the Kayama et al. dataset, 72 pairs of PCR primers with lengths ranging from 19 to 22 base pairs were designed with Primer3 software to amplify 31 double-stranded templates, with lengths ranging from 435 to 481 bases. Templates were synthetically generated using the OE-PCR technique for partial 16S rRNA nucleotide sequences (v6-v8). The design of primers was based on their specificity for the corresponding templates, targeting an amplification length of approximately 100-150 base pairs. It is assumed that primer-template binding is not limited to the primer’s full length and only a part of the 3’ end may bind.

In contrast to the Döring et al. dataset, the Kayama et al. dataset does not provide direct documentation of various experimental features. The data available in their study includes only the sequences of the template and primers, along with the corresponding amplification results. Nevertheless, features such as Gibbs free energy and mismatch information can be analyzed solely based on the primer and template sequences.

### 2.2. Feature Selection

Given the diversity of parameters across the PCR experiment, it is crucial to identify the features that have the greatest impact on PCR amplification outcomes. To achieve this goal, we applied a feature selection method to the Döring et al. dataset (Figure 1-b). Due to the common nature of the problem, we can select the most important features in this dataset and then analyze these features in the dataset of Kayama et al. to perform prediction using our proposed model.

We used an embedded method to identify features that significantly influenced PCR results. In this method, the subset of selected features achieves the best accuracy when derived during the construction of a classifier [25]. Embedded methods leverage each iteration of the model training process for feature selection, effectively identifying features that provide the greatest benefit for enhancing data training. During the training phase, the classifier modifies its internal parameters and assesses the appropriate weights for each feature in order to achieve the best classification accuracy [26]. This can be achieved using various classifiers, such as the random forest classifier used in our study.

By applying this algorithm to the dataset, we identified several key features that are most important for predicting PCR amplification, shown in Figure 2. As we can see, Gibbs free energy is the most important feature that specifies the success of binding. Although primer efficiency is another important feature, it was evaluated during the initial design using Primer3 and does not require further consideration. Other important features in determining the outcome are based on match and mismatch information, such as their number and location, which directly influence the stability of the binding, and thus the amplification outcome. In the following sections, we will propose our approach to analyse Gibbs free energy and match and mismatch information.

**Figure 2:**
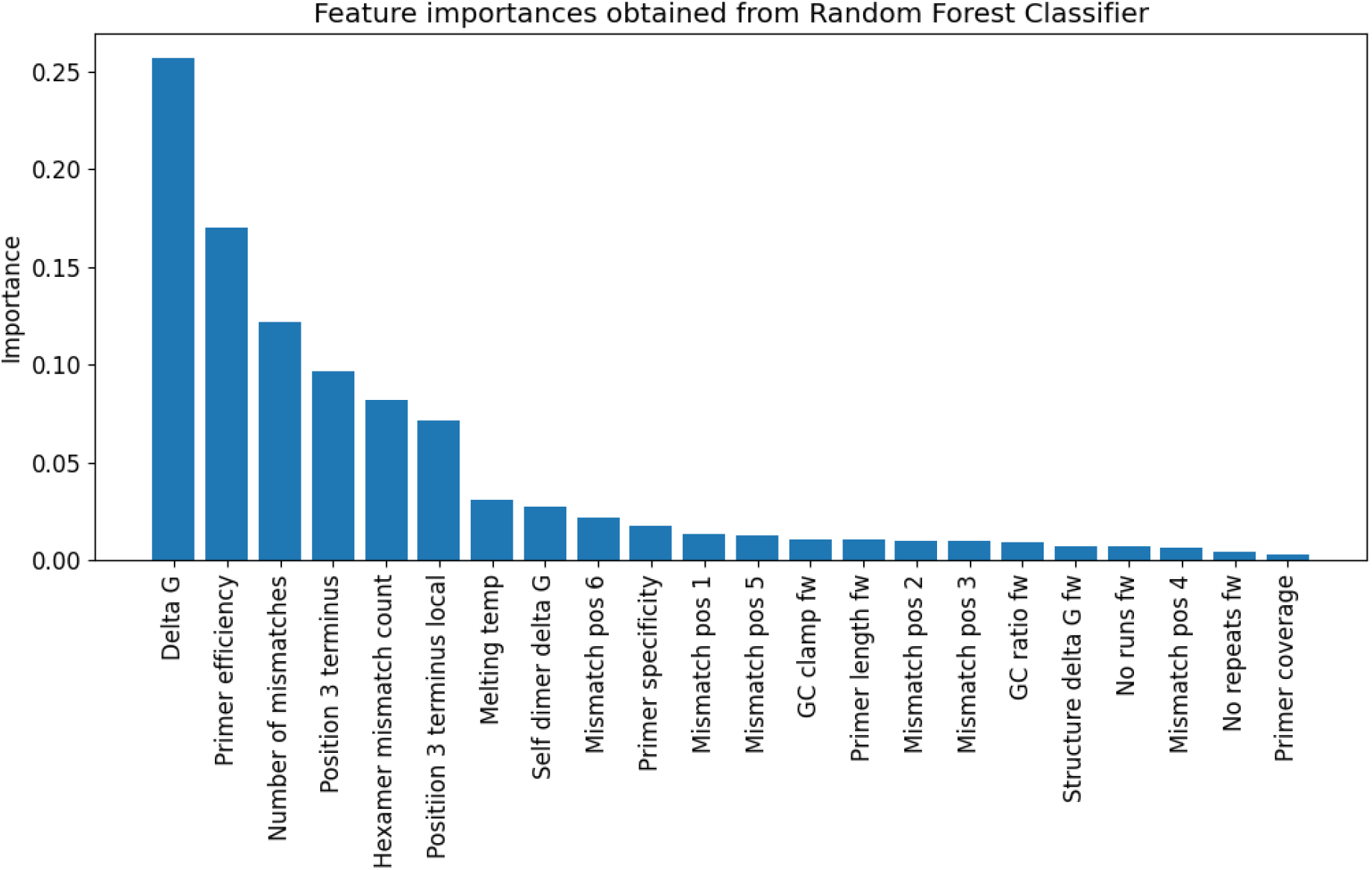
Results of feature selection using Random Forest Classifier on the first dataset. We chose the first six features as the most important features for predicting PCR amplification results.

### 2.3. Binding site identification

Primers can bind to various positions on the template, and the stability of these bindings is determined by the Gibbs free energy (Δ*G*). The lower the Δ*G*, the more stable the binding. Because the complementary region is short enough, the most stable pairing is likely to be the priming position (Figure 1-c). To identify the most stable binding position for each primer-template pair, we should slide the primer on the template and calculate Δ*G* for each position of binding and choose the place with minimum Δ*G*, as shown in Figure 3. Δ*G* is defined as below,

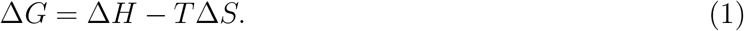

**Figure 3:**
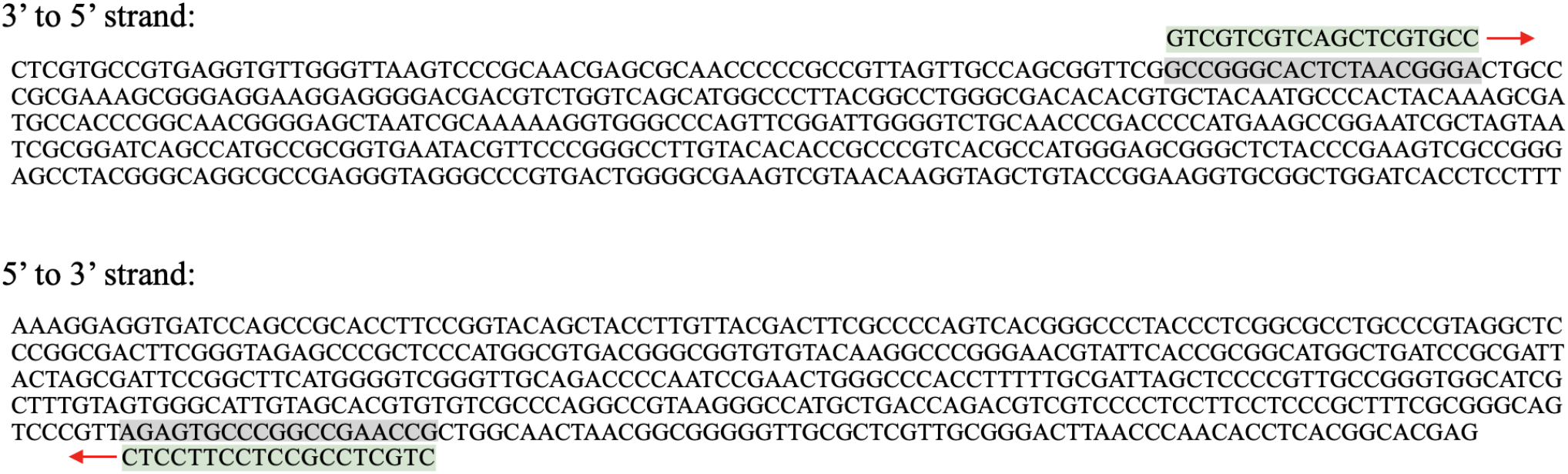
Sliding forward and reverse primer on the corresponding template strand and calculating the Δ*G* of each place. The place withthe minimum Δ*G* may be the binding position.

In this formula, Δ*H* and Δ*S* are enthalpy and entropy of sequence, respectively. T is the annealing temperature, which is 56*^◦^* for this experiment. We can calculate Δ*H* and Δ*S* using the approach presented in the study of Horne et al. [27]. In this approach for a pair of primer-template, Δ*H* is calculated by adding up the enthalpy of each pair of neighbouring bases as shown below,

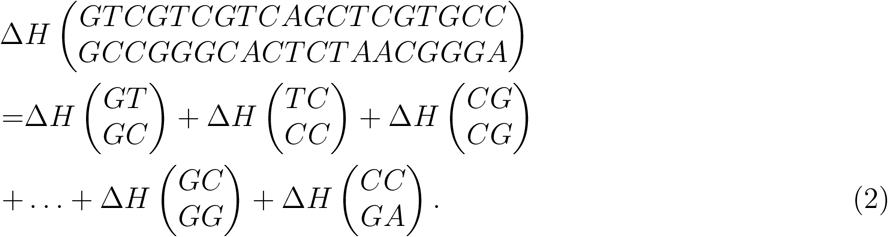

Δ*S* can be calculated using the same way. Δ*H* and Δ*S* for each type of neighbouring base pairs are proposed in the same study. We calculate Δ*G* using this method for both forward and reverse primers. After this calculation for all of the possible binding positions for each primer, we choose places for binding positions of forward and reverse primers with minimum Δ*G* that are at least 100 base pairs apart.

### 2.4. Labelling Primer-Template Matches and Data Augmentation

In Section 2.2, we concluded that Δ*G* and match/mismatch information are the most influential features for predicting PCR amplification. After determining the binding site using Δ*G* in Section 2.3, the next step is to characterize match and mismatch positions in the primer-template pair.

Since the primer binding site can be shorter than the primer length, matches or mismatches may also occur in nucleotides preceding or following the main binding site. Therefore, each binding position must be represented in a way that distinguishes matched and mismatched regions both at and around the binding site (Figure 1-d).

To encode this information, we assigned specific words to represent different types of binding events. Using cosine similarity analysis of words close to the word “matched” and “mismatched” in BERT (Figure 5), we identified the words “correspondence,” “pass,” “equal,” and “accordance” as semantically close and thus suitable for match positions, while “disparateness,” “incongruity,” “incomparability,” and “discrepancy” were found to cluster together and were therefore selected for mismatch positions. Figure 5 illustrates how words within each group exhibit strong internal similarity but remain clearly distinct from those in the opposite group.

For each base pair in the forward and reverse primers, we assigned a word according to its class (match or mismatch), type (forward or reverse), and position (within or outside the binding site). The forward and reverse sequences were then concatenated to form a single representative sequence for each primer-template pair (Figure 4). This process was repeated for all primer-template combinations, resulting in a labeled dataset that systematically captures binding specificity.

**Figure 4:**
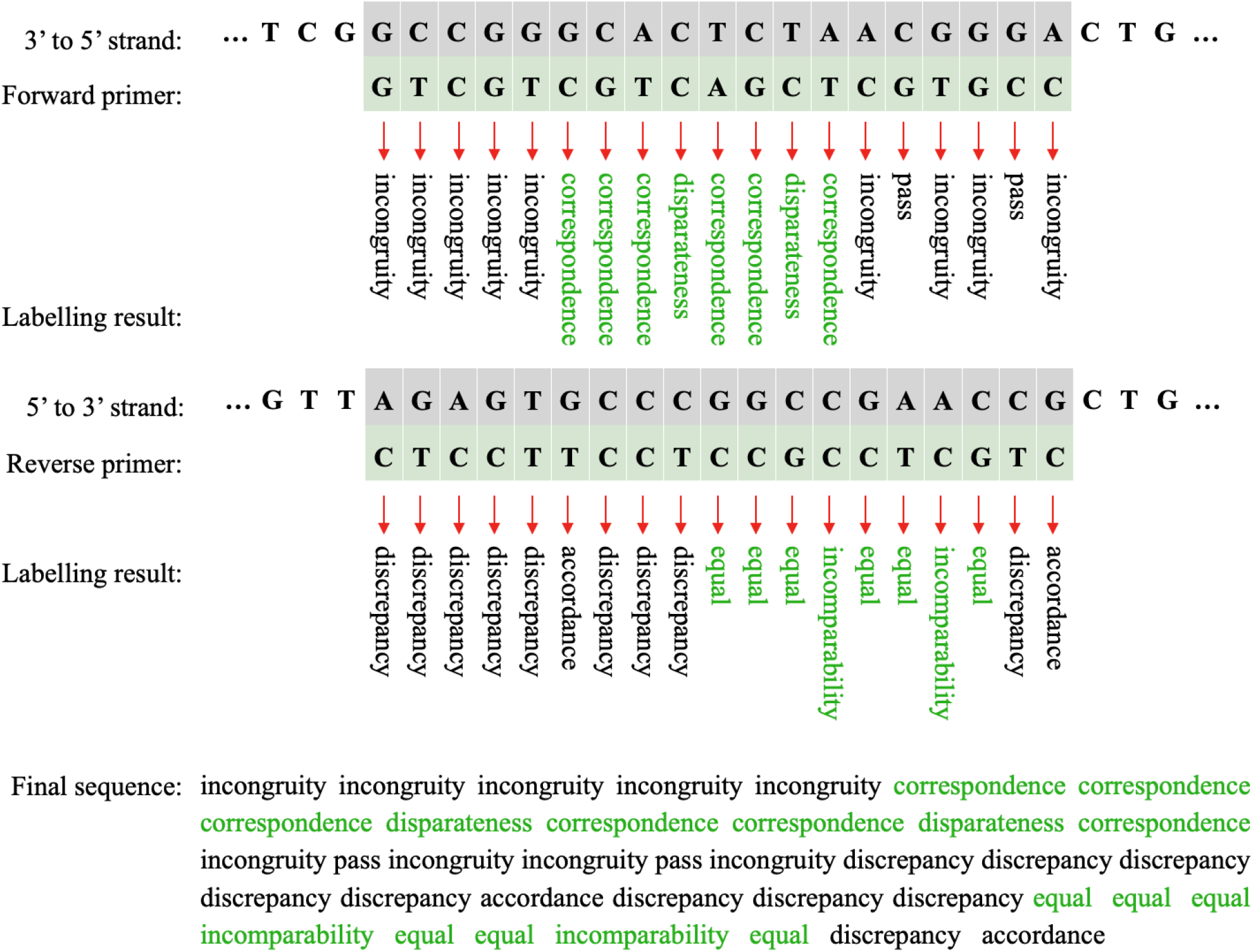
Primer-template labelling based on matches and mismatches in each position. Words are assigned based on match/mismatch status, primer type (forward or reverse), and position (within or outside the binding site). Green words indicate the binding site and black words indicate regions before or after it. The final sequence is obtained by concatenating forward and reverse primer labelling results.

**Figure 5:**
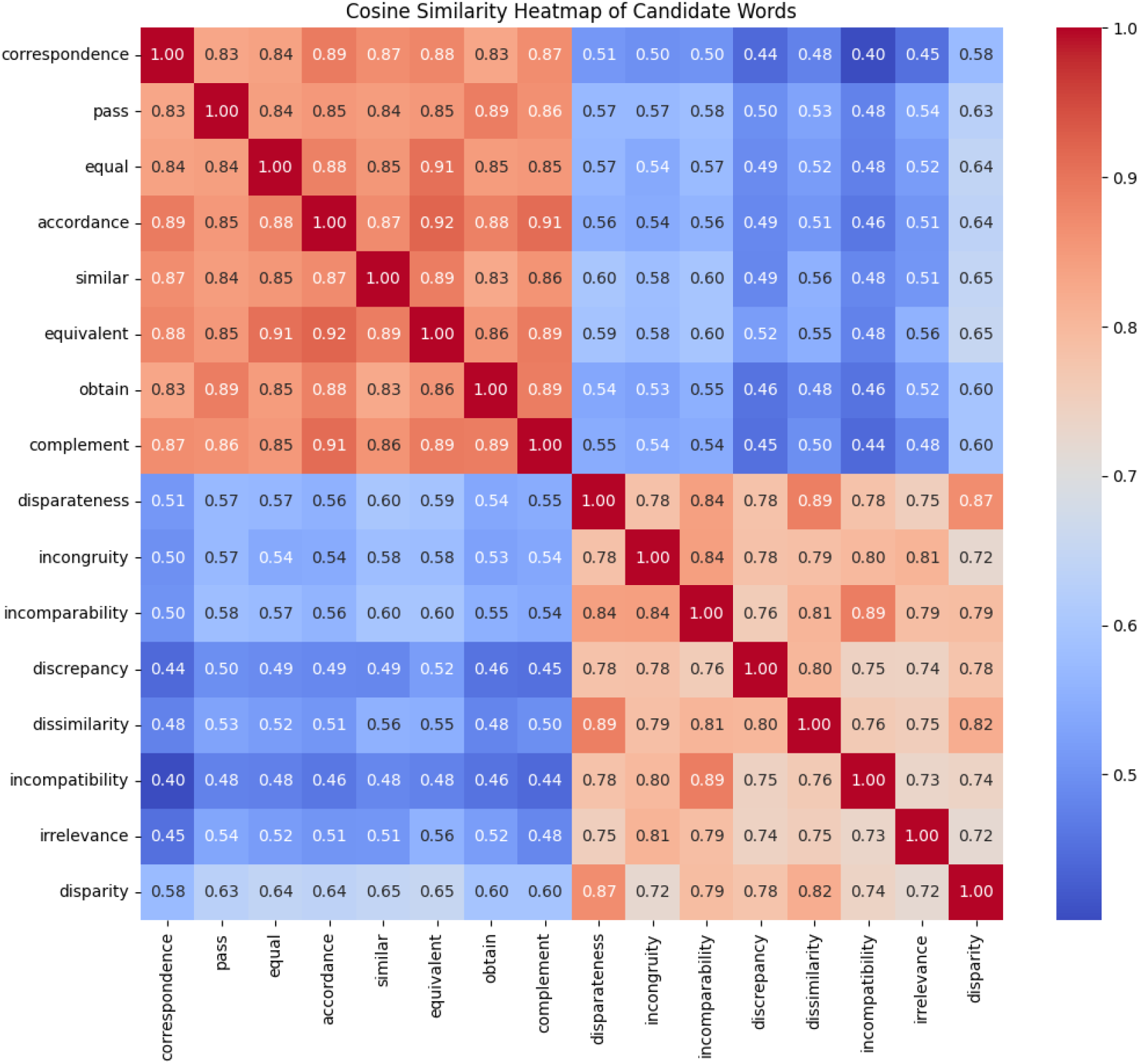
Cosine similarities among words used to label match and mismatch positions. Words within each group (match or mismatch) exhibit strong semantic similarity, while inter-group similarities remain low, highlighting clear separation between the two categories.

Because the dataset size was relatively small, we applied a data augmentation strategy strictly in training data to improve model generalization [28]. To completely prevent fold-level data leakage, all dataset splitting for cross-validation was performed prior to augmentation. The validation and holdout test sets contained only original, unaugmented sequences. Specifically, we created eight augmented copies of the dataset, each incorporating one targeted synonym replacement: “correspondence” *→* “similar”, “disparateness” *→* “dissimilarity”, “pass” *→* “obtain”, “incongruity” *→* “inconvenient”, “equal” *→* “equivalent”, “incomparability” *→* “incompatibility”, “accordance” *→* “complement”, and “discrepancy” *→* “disparity”. The complete mapping of assigned words for each category and position is shown in Table 1. This augmentation preserved the semantic relationships among words while enriching the diversity of the training samples.

**Table 1:** Assigned words for each primer-template binding type in the augmented dataset.

|  |  | **Corresponding label** |  |
|---|---|---|---|
| **Primer** |  | **At the binding site** | **Before or after the binding site** |
| Forward | Match | correspondence<br>similar | pass<br>obtain |
|  | Mismatch | disparateness<br>dissimilarity | incongruity<br>inconvenient |
| Reverse | Match | equal<br>equivalent | accordance<br>complement |
|  | Mismatch | incomparability<br>incompatibility | discrepancy<br>disparity |

### 2.5. BERT Tokenizer

We used a BERT tokenizer to tokenize each sequence by its words (Figure 1-f). BERT tokenization transforms text into tokens that represent individual words or subwords, allowing the model to understand and process the text effectively [24]. This tokenization captures contextual relationships and semantic meanings, which are crucial for tasks such as text classification or sequence prediction in models like BiLSTM. We use this tokenization to capture patterns of match and mismatches in the primer-template pairs of PCR.

### 2.6. The Proposed Model

To model the complex relationships between mismatches and matches in primer-template sequences, we adopted a multi-input, multi-branch architecture. The core of this model, responsible for processing the sequential data, utilizes a Convolutional Neural Network (CNN) layer followed by a Bidirectional Long Short-Term Memory (BiLSTM) network. This design choice was motivated by the sequential and contextual nature of our modeled primer-template interactions in PCR.

Recurrent Neural Networks (RNNs) are a class of models well-suited for sequential data. However, standard RNNs are limited in their ability to capture long-range dependencies due to the vanishing gradient problem. LSTM networks address this issue by incorporating gating mechanisms that allow them to preserve information over longer sequences. While LSTMs improve upon RNNs, they typically operate in a unidirectional manner which restricts their ability to capture full contextual information.

In our task, understanding the outcome of a primer-template match requires capturing patterns that span both forward and reverse interactions. This is especially critical because amplification can be influenced by interactions that occur on either sides of the primer-template interaction. A mismatch may have different effects depending on its position relative to both ends of the primer and on whether it’s part of the forward or reverse binding process. For instance, a mismatch near the 3’ end of a primer often has a greater negative impact on amplification than one near the 5’ end. The bidirectional architecture enables the model to learn such nuanced patterns across the entire sequence. Therefore, we employed a BiLSTM, which reads the sequence in both directions and can effectively learn dependencies across the entire primer-template region from both orientations. Compared to traditional machine learning approaches, BiLSTM-based models like Enhancer-LSTMAtt [29] offer better performance by capturing long-range dependencies in DNA sequences.

Before feeding the sequence to the BiLSTM, we first process it with a 1-dimentional Convolutional Neural Network (CNN) layer. While the BiLSTM is excellent at capturing long-range or global dependencies, the CNN excels at identifying local patterns. In our sequence, a “local pattern” might be a specific three-word phrase like ‘pass incongruity pass’ or ‘incomparability incomparability equal’. The CNN layer acts as a local feature extractor, scanning the sequence for these small, significant motifs. By applying a CNN layer to the BERT embeddings, the model learns to recognize these important local n-grams. The output of this layer—a set of feature maps highlighting local patterns—is then passed to the BiLSTM, which can then learn the sequential order and long-range relationships between these identified local features.

A key strength of our model is its multi-input, multi-branch nature, which allows it to synthesize information from disparate but complementary sources. Instead of relying solely on the text sequence, the model integrates two distinct types of data:

- Sequential Interaction Data (BERT-CNN-BiLSTM): This is the primary branch, as described above. It begins by encoding the sequence of interaction words (e.g., ‘pass’, ‘discrepancy’) using a pre-trained bert-base-cased model. This provides rich, contextual embeddings (768 dimensions) for each word. This BERT-derived sequence is then processed by the CNN and BiLSTM layers to understand the complex grammar and long-range dependencies of the primer-template interactions.
- Hand-Crafted Biological Features: We recognized that certain high-level biological metrics are strong predictors of PCR success. This branch accepts a vector of 6 engineered features, including the GC content of the forward, reverse, and template sequences, the total mismatch count, the maximum continuous mismatch count, and metrics for 3-prime hexamer mismatches. These features are processed through their own small feed-forward network, allowing the model to learn non-linear relationships between these explicit biological rules.

These branches are then concatenated into a single, comprehensive feature vector. This combined vector is passed through a final classifier populated with ReLU activation functions, allowing the model to make its final prediction by weighing evidence from the deep-learned sequence patterns, the explicit biological rules, and the latent primer-specific properties. This hybrid approach is powerful because it grounds the abstract representations learned by the BiLSTM with concrete, domain-specific biological knowledge.

Key hyperparameters for the model were optimized and set as follows. We ran the model for 40 epochs. The ‘bert-base-cased’ model was made trainable to allow it to fine-tune its word embeddings to our specific domain. The sequence branch uses a CNN layer with 16 filters and a kernel size of 3, followed by a BiLSTM with 16 units. Using a relatively small number of units in both the CNN and LSTM layers helps to prevent overfitting on this specialized dataset.

The model is trained using the Adam optimizer, chosen for its adaptive learning rate properties, with an initial learning rate of 3e-4 and a weight-decay coefficient of 1e-3. The learning rate was reduced by a factor of 0.5 when PR-AUC did not improve for 5 consecutive epochs, with a minimum learning rate of 1e-6. We employ a batch size of 64. To combat overfitting, two Dropout layers are used with a rate of 0.5. The model is compiled with binary cross-entropy loss, as this is a binary classification task. Finally, class weights are computed and applied during training to counteract the imbalanced nature of the dataset, ensuring the model does not simply favor the majority class.

## 3. Results

To evaluate model performance, we used grouped 5-fold cross-validation on the development dataset, with each model trained for up to 40 epochs. The best epoch in each fold was selected according to the highest inner-validation PR-AUC. The model achieved an accuracy of 0.838 ± 0.010, a sensitivity of 0.716 ± 0.099, a specificity of 0.877 ± 0.033, and an AUC of 0.878 ± 0.033 across the five outer validation folds. Figure 6 presents the training and validation accuracy curves, while Figure 7 shows the corresponding ROC curves.

**Figure 6:**
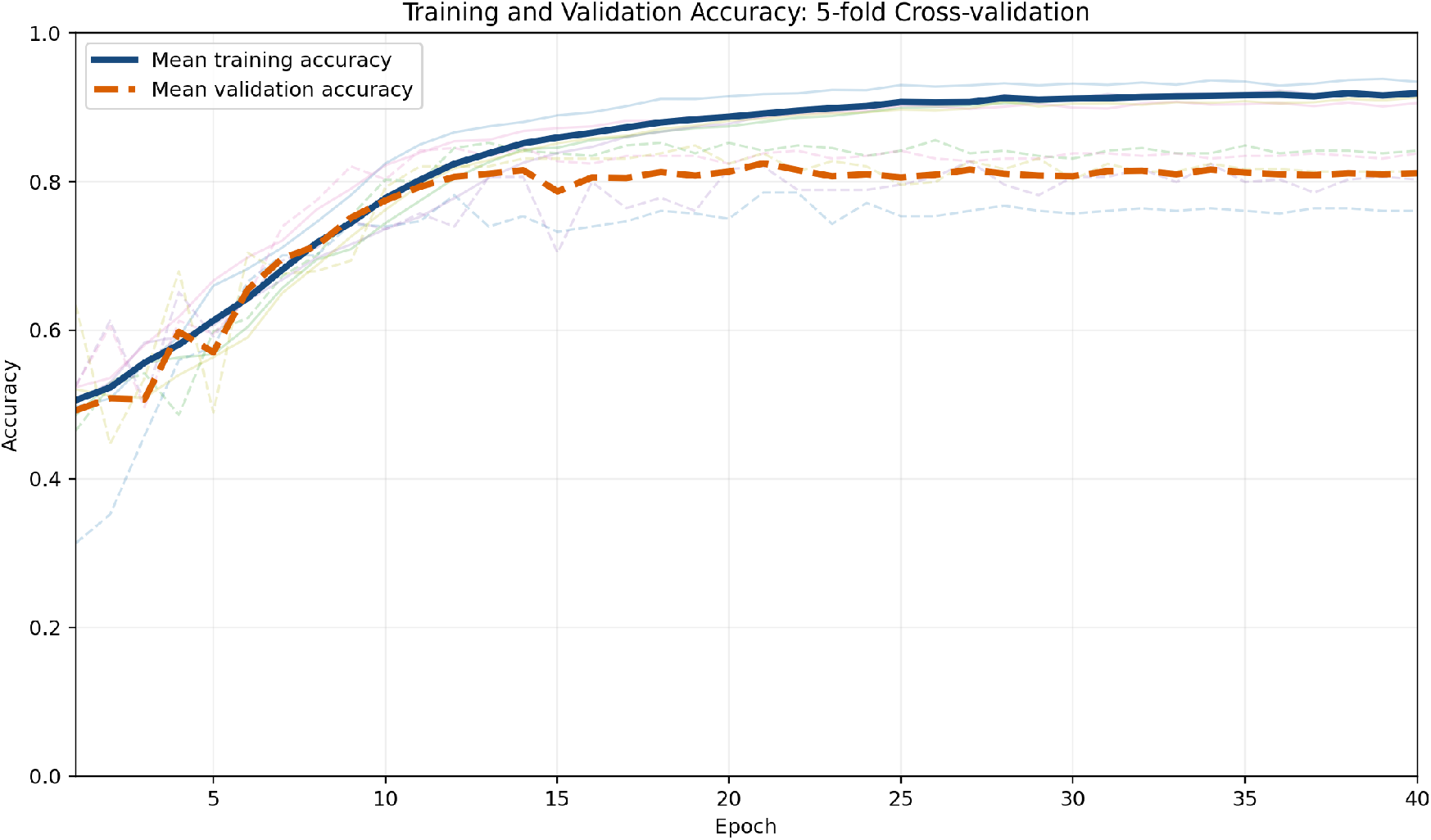
Training and Validation accuracy results for 5-fold cross-validation.

**Figure 7:**
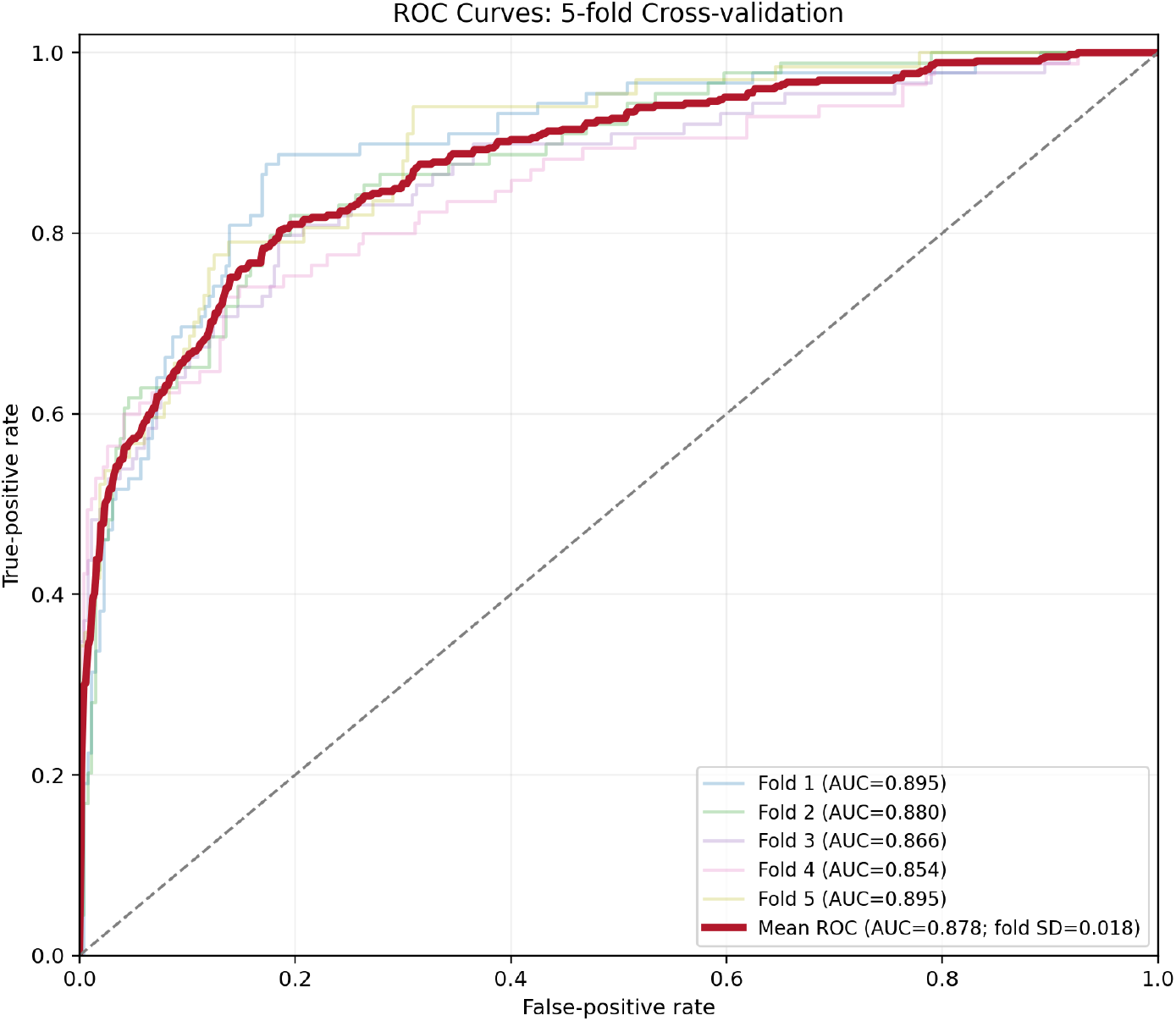
ROC curve for 5-fold cross-validation.

Before model training, 20% of the complete dataset was reserved as an untouched final test set and excluded. The final model was trained using the median of the best epochs selected across the five folds. On the untouched final test set, it achieved an accuracy of 0.859, a sensitivity of 0.732, a specificity of 0.904, and an AUC of 0.900.

## 4. Discussion

In the raw data that we have in the PCR experiment, we only have primers and template sequences. If we want to predict the outcome, we need a model that can learn base-pairing rules, the significance of a match versus a mismatch, and the positional information as we know that the position of mismatches can effect the PCR amplification result. A conventional nucleotide-based model must first learn the fundamental rules of base-pairing from scratch before it can begin to understand the higher-order patterns that govern PCR amplification.

In our proposed model, We presented a new labelling strategy based on the location and number of mismatches in the forward and reverse primer-template binding. We claim that when the aim is to identify the binding pattern of the primer to the template, the specific types of nucleotides involved are not critical. What matters is whether the paired nucleotides are matches or mismatches. For instance, there is no distinction between C-G and A-T connections, as both are considered matches. Similarly, T-T and A-A connections are both mismatches and in our proposed model, there is no need to assign separate codes for each. The type of bases present in a nucleotide connection only affects the calculation of Gibbs free energy since each consecutive pair of nucleotides has its own enthalpy and entropy values. In our proposed model, the calculation of Gibbs free energy occurs before word formation. After selecting the appropriate location for primer binding, the specific nucleotide types are no longer necessary. It suffices to know whether the connection is a match or a mismatch. In our semantic labeling strategy, we reframe the biological problem of primer-template interaction into a linguistic context, thereby enabling the use of powerful transfer learning from pre-trained language models. We represented matches and mismatches using two distinct groups of English words that are semantically related to the concepts of “match” and “mismatch.” The advantage of this complex encoding is the creation of a richer, more nuanced feature space. By leveraging a pre-trained BERT tokenizer, we are providing our model with high-dimensional embeddings that are already infused with a deep, pre-existing understanding of similarity and opposition.

To validate our proposed multi-input, multi-branch architecture and quantify the contribution of its key components, we conducted an ablation study. We established a baseline model that represents the architectural backbone of our sequence processing branch but is ablated of all advanced and domain-specific inputs. This baseline model consisted of a CNN layer followed by a BiLSTM, similar to our main model. However, it critically differed in the following ways: 1) Instead of using fine-tuned, 768-dimension BERT embeddings, this model used a standard Keras Embedding layer trained from scratch on our vocabulary. 2) Single-Input Model: It was a single-input model that did not incorporate the multi-branch design. The hand-crafted biological features (e.g., GC content, mismatch counts) were completely removed. 3) No Data Augmentation: The model was trained only on the original, un-augmented dataset. This simplified baseline model was trained and evaluated using the same 5-fold cross-validation methodology as our full model to ensure a fair comparison. The performance of this baseline model was substantially lower than our full, multi-branch architecture. The aggregated 5-fold cross-validation results yielded an average accuracy of 0.630 ± 0.031% and an AUC of 0.673 ± 0.014. The stark contrast between these metrics and those achieved by our final, complex model provides a clear justification for our architectural choices. It demonstrates that a simple CNN-BiLSTM, while capable of learning some basic patterns from the interaction sequence, is insufficient for this complex classification task. The results strongly suggest that the significant performance lift of our final model is driven by the synergistic combination of the rich, contextual representations from the fine-tuned BERT model, and the explicit domain knowledge injected via the biological feature branch.

We further evaluated the contributions of semantic data augmentation and cosine-similarity-based word selection through two additional ablation experiments, while keeping the model architecture, biological feature branch, training settings, and grouped 5-fold cross-validation procedure unchanged. First, we evaluated the necessity of our specific vocabulary. By deliberately selecting English words that are semantically clustered—grouping synonyms for “match” strictly separate from those for “mismatch”—we translated the physical topology of primer-template binding into a coherent linguistic framework. This semantic proximity is what allows the pre-trained BERT model to intuitively distinguish biological states based on word relationships. When we replaced these cosine-similarity-selected words with randomly chosen English words, the model’s accuracy dropped significantly to 0.721 ± 0.037, and the AUC fell to 0.806 ± 0.016. This sharp decline demonstrates that simply assigning arbitrary tokens is insufficient; mapping structural topologies to semantically meaningful words is vital for enabling BERT’s pre-trained attention mechanisms to recognize complex grammatical mismatch patterns. Second, we assessed the role of our semantic data augmentation. Because small datasets in computational biology frequently lead to model overfitting, we introduced synonym replacement to synthetically expand the training data without altering the underlying biological truth. To quantify its necessity, we trained the model completely devoid of augmentation, which resulted in a reduced accuracy of 0.788 ± 0.062 and an AUC of 0.839 ± 0.025. This drop confirms that our semantic augmentation strategy effectively enriches the linguistic diversity of the training space, acting as a crucial component to prevent overfitting and ensure robust generalization.

A direct comparison of model performance with prior works is crucial for benchmarking. We can compare our model with the studies of Kayama et al. [19] and Bai et al. [20], as we used the same dataset as them. In their studies, the coded sequence consists of 5-letter words formed from 25 characters of the English alphabet, utilizing both lowercase and uppercase letters, dependent on the position and type of nucleotide pairs. The characters used correspond to each type of connection between nucleotides, with every five consecutive characters forming a word. Moreover, to emphasize features like closeness to the 3’ terminal for the RNN model, their models repeat words based on their positional placement. In contrast, uses meaningful English words instead of single letters. Our proposed model does not require such repetition of words. This is because it labels connections between primers and templates based solely on matches and mismatches. As a result, identifying all relevant features becomes more straightforward, especially when we utilize a BERT tokenizer to recognize patterns in the sequence, considering their type and position, following a CNN to find local patterns.

Comparing different models sometimes is complicated by differences in data partitioning. Since we did not have access to the original, segregated test sets used in Kayama et al. and Bai et al., we established our own rigorous evaluation framework. We first partitioned the entire available dataset into an 80% training/validation set and a 20% holdout test set. All cross-validation metrics reported for our model were generated exclusively from the 80% partition, while the 20% set was reserved for a final, unseen test. The mentioned prior works appear to have used the entirety of this same dataset for their reported cross-validation. Therefore, the most equitable comparison is between our 5-fold cross-validation metrics (derived from our 80% training/validation set) and the validation metrics reported in those studies. A summary of this comparison is presented in Table 2. Our proposed multi-input model demonstrates a significant improvement in overall performance and a much more balanced predictive profile.

**Table 2:** Comparing results with prior studies.

| **Reference** | **Method** | **Sensitivity** | **Specificity** | **Accuracy** | AUC |
|---|---|---|---|---|---|
| Kayama et al [19] | LSTM | 0.563 ± 0.062 | 0.882 ± 0.024 | 0.803 ± 0.029 | - |
| Bai et al [20] | Attention-BiLSTM | 0.570±0.06 | **0.905±0.03** | 0.822±0.02 | 0.80 |
| Current study | CNN-BiLSTM | **0.710±0.089** | 0.879±0.034 | **0.838±0.010** | **0.88** |

Our model achieved the highest mean accuracy of 83.8%, surpassing both Bai et al. (82.2%) and Kayama et al. (80.3%).The most notable advancement is in sensitivity. Our model’s sensitivity of 71.6% is dramatically higher than that of Bai et al. (57.0%) and of Kayama et al. (56.3%). This indicates that our model is vastly superior at correctly identifying true positive cases (successful PCR outcomes), which is a critical weakness in previous models. While the models from Bai et al. and Kayama et al. report slightly higher specificity (90.5% and 88.2%, respectively), our model maintains a highly competitive specificity of 87.7%. This small trade-off is exceptionally favorable, as we gain over 14 percentage points in sensitivity (vs. Bai et al.) while conceding only 3 points in specificity. This demonstrates that our model is far more balanced and does not suffer from the heavy bias towards the majority negative class (failed PCR outcomes) that seemingly affects the other models. Furthermore, our model exhibits greater stability. The standard deviations for all our key metrics (e.g., ± 0.010 for accuracy) are substantially lower than those reported in the other studies. This suggests that our model’s performance is more consistent and less dependent on the specific data fold, indicating robust generalization. Finally, our model achieved a mean AUC of 0.87.8%. This high AUC in comparison with reported AUC in Bai et al. study (80% with high variance in folds), coupled with the corresponding aggregated ROC curve, confirms the model’s superior discriminative power across all decision thresholds, solidifying its status as a more reliable and robust classifier for this complex biological problem.

In addition to comparing our results with studies that used the same dataset, we reviewed other relevant works that relied on a predefined table of features, predicting the outcome with machine learning models. Cordaro et al. [12] and Döring et al. [18] have achieved accuracies of 81% and 95%. Kronenberger et al. [17] yielded 92.4% and 96.5% for two different types of probes in their study. The feature-based approaches require comprehensive documentation of all relevant features, which can be challenging, especially as new data characteristics emerge. In contrast, our method circumvents this limitation by predicting PCR outcomes directly from the template and primer sequences, without the need for extensive feature documentation. This streamlined approach not only simplifies the predictive process but also enhances adaptability to new data, making it more efficient and scalable.

Furthermore, the lack of augmentation in all of the prior works while having small datasets, limits the robustness and generalizability of their models, as small datasets can lead to overfitting. In contrast, our study incorporates augmentation, which improves model performance and reliability, especially in low-data settings.

Additionally, while established software tools like Primer3 are essential for the initial design of candidate primers using thermodynamic heuristics, they are not designed to reliably predict the final binary amplification outcome when complex mismatch patterns occur. Our proposed model serves as an in silico bridge between this initial design phase and physical experimentation. By acting as a pre-synthesis filter, it shifts the burden of troubleshooting from the physical laboratory—where iterative re-design and re-testing are costly and slow—to a rapid computational step. This provides a distinct practical advantage, particularly for high-throughput applications such as multiplex PCR or large-scale next-generation sequencing (NGS) library preparation, where minimizing the experimental failure rate is critical. However, we acknowledge that while our current metrics demonstrate strong algorithmic improvements, particularly in sensitivity, a comprehensive real-world wet-lab benchmarking study to quantify the exact economic and temporal savings compared to traditional laboratory troubleshooting remains a necessary future endeavor.

In this study, model training and validation are centered on a specific set of previously published templates and experimental settings. While we ensured rigorous internal validation by isolating a strict 20% holdout test set prior to any augmentation or training, the evaluation of the model’s broad generalizability can be further expanded through the use of fully independent, external test sets. Because the current training data may not encompass the massive variability seen in different PCR applications (e.g., highly multiplexed assays, extreme GC-rich targets, or diverse environmental DNA samples), its applicability to sequence characteristics and experimental conditions outside the scope of this dataset cannot be guaranteed. At this stage, our framework serves as a proof-of-concept demonstrating the powerful potential of NLP-based semantic tokenization for PCR prediction. Generating a proprietary, large-scale dataset to encompass a wider variety of failure patterns remains prohibitively expensive and time-consuming for a purely computational study. Consequently, comprehensive benchmarking against independent, diverse wet-lab datasets is a necessary future endeavor before this model can be broadly deployed across varied experimental PCR settings.

## 5. Conclusion

In this study, we predicted PCR amplification by modeling primer-template interactions, using BERT tokenization, and a multi-input, multi-branch deep learning model. First, we identified the most important features for the amplification result prediction based on a dataset containing different features, which resulted in Δ*G*, primer efficiency, and some other features related to match and mismatch positions. By analyzing each binding place between each pair of primer and template in the dataset, we found the place with the least Δ*G*, which is the expected binding site. The primer-template binding was then modeled as a sequence by assigning a word to each match and mismatches at the binding site or immediately before and after it. Different words were assigned to represent matches and mismatches at the binding site and its adjacent positions, resulting in a unique sequence for each primer-template combination.

We then used a BERT tokenizer to tokenize each sequence based on its words. This collection of encoded sequences formed the primary input branch for a CNN BiLSTM, which was integrated with the other branch containing biological features. By running the full model on this combined input data with 5-fold cross-validation, an accuracy of 0.838±0.010, a sensitivity of 0.716±0.099, a specificity of 0.877±0.033, and an AUC of 0.878±0.018was achieved. On the untouched final holdout test set, the model achieved an accuracy of 0.859, a sensitivity of 0.732, and a specificity of 0.9044, and an AUC of 0.900. Our presented model not only has higher accuracy, sensitivity, and AUC, but is also simpler than the models presented in previous studies by proposing more meaningful and applicable labelling. This approach enables predictions of amplification results based solely on template and primer sequences, thereby saving time and costs associated with conducting PCR experiments.

## 6. CRediT authorship contribution statement

Niloofar Latifian: Data curation, Formal analysis, Investigation, Methodology, Visualization, Writing – original draft, Writing – review and editing.

Naghme Nazer: Conceptualization, Methodology, Project administration, Writing – review and editing.

Amir Masoud Jafarpisheh: Investigation, Methodology, Writing – review and editing. Babak Khalaj: Project administration, Supervision, Writing – review and editing.

## 7. Data and Code Availability

The data and code supporting this work are openly available at: github.com/niloofarlatifian/PCR-Amplification-Prediction.

## 8. Declaration of competing interest

The authors have no conflict of interest to declare.

## 9. Acknowledgment

We thank Professor Daiji Endoh at the Department of Radiation Biology, School of Veterinary Medicine, Rakuno Gakuen University for kindly providing the data used in this study.

## 10. Funding Sources

This research did not receive any specific grant from funding agencies in the public, commercial, or not-for-profit sectors.

## 11. Declaration of generative AI and AI-assisted technologies in the writing process

During the preparation of this work, the authors used ChatGPT and Gemini in order to improve language and readability. After using this tool, the authors reviewed and edited the content as needed and take full responsibility for the content of the publication.

## Notes

https://github.com/niloofarlatifian/PCR-Amplification-Prediction

### Competing Interest Statement

The authors have declared no competing interest.

### Summary of Updates

Removed ID Embeddings: Completely removed primer and template ID embeddings and retrained the model using only BERT sequence data and biological features so we are sure that it learns transferable PCR rules rather than dataset-specific artifacts. Adjusted Regularization & Explained Encoding: Modified the Dropout rate so that training accuracy properly tracks higher than validation accuracy (updated in Figure 6), confirmed no data leakage occurred during cross-validation splits, and added biological justification for mapping nucleotide structures to English semantic tokens. Tempered Generalizability Claims: Clarified the use of a strictly isolated 20% holdout test set, and revised the Discussion to note that broader external validation is a necessary next step. Added Targeted Ablation Studies: Added two controlled ablation experiments isolating the impact of the semantic encoding (replacing targeted words with random ones dropped accuracy to 72.1%) and semantic augmentation (removing it reduced accuracy to 78.8%). Clarified Practical Value: Expanded the Discussion to better explain how the model serves as an in silico pre-synthesis quality-control filter (especially for high-throughput tasks like NGS) and highlighted its 71.6% sensitivity in reducing false negatives compared to traditional primer-design tools.

